# Non-native Interactions Explain the Folding Rate Differences in *α*-Spectrin Domains and the Origin of Internal Friction Effects

**DOI:** 10.1101/232116

**Authors:** Fernando Bruno da Silva, Vinícius G. Contessoto, Vinícius M. de Oliveira, Jane Clarke, Vitor B. P. Leite

## Abstract

Recent experimental and computational studies have shown the influence of internal friction in protein folding dynamics. However, uncertainty remains over its molecular origin. α-spectrin experimental results indicate that R15 domain folds three orders of magnitude faster than its homologous R16 and R17. Such anomalous observations are usually attributed to the influence of internal friction on protein folding rates. To study this phenomenon, we carried out molecular dynamics simulations with structure-based Cα models, in which the folding process of *α*-spectrin domains was investigated by adding non-native interactions. The simulations take into account the hydrophobic and the electrostatic contributions separately. The folding time results have shown a qualitative agreement with experimental data. We have also investigated mutations in R16 and R17, and the simulation folding time results correlate with the observed experimental ones. We suggest that the origin of the internal friction emerges from a cooperativity effect of these non-native interactions.

## Introduction

The concept of internal friction in the folding processes has been extensively studied by experimental and computational groups.^1–7^ Some experimental measurements have identified a deviation in the expected relationship between the folding rate and solvent viscosity.^1,8^ The suggestion is that the folding process is also influenced by internal collisions within the protein, which could explain low viscosity dependence of the folding rates.^1^ The internal friction generated by collisions within the protein may be interpreted as an energy dissipation mechanism that does not contribute to return the protein to its native state in the folding process.^8,9^ In addition, the internal friction in the folding process may be related to roughness in the energy landscape^10,11^ and it may also be associated with secondary structure mis-docking induced by non-native interactions.^4,12^ The folding time deviation related to internal friction is less evident for certain protein motifs, such as all-*β* and *α/β* proteins.^13,14^ On the other hand, there is a higher folding time deviation for zero viscosity extrapolation in *α*-helical proteins, indicating that they may be more affected by internal friction.^12,15–17^

Recently, three specific domains from *α*-spectrin were found to display differences in their folding rates, and it was suggested that these differences might be related to internal friction.^10–12,18–21^ The R15, R16, and R17 domains of *α*-spectrin are composed of an elongated three helix bundle and display similar structures, thermodynamic stabilities and *β*-Tanford values.^19,22–24^ Clarke and co-workers reported that R15 folds and unfolds three orders of magnitude faster than its homologues R16 and R17.^19^ A significant fraction of this difference in folding rate could be ascribed directly to differences in internal friction. Early computational efforts show a good agreement of the Φ-values with experiments,^4^ but the reason for the differences in folding rates and the origins of internal friction remain unclear.

In the present study, the molecular dynamics simulation was carried out using the Structure-Based C_*α*_ Model (SBM-C_*α*_) with the addition of non-native interactions, which take into account the hydrophobic and electrostatic contributions. In order to describe the folding process of R15, R16, and R17, we performed four types of simulation: (i) SBM-C_*α*_ without non-native interactions, (ii) SBM-C_*α*_ with hydrophobic non-native contribution, (iii) SBM-C_*α*_ with electrostatic non-native contribution and, (iv) SBM-C_*α*_ with both non-native contributions. Folding time calculation was performed for each model and compared with the experimental data. The mutations that speed up or slow down experimental folding time were analyzed computationally and the results present a qualitative agreement. Free energy profiles and folding routes were also calculated, and the results corroborated the kinetic results, permitting discussion of internal friction effects in the R15, R16 and R17 domains in terms of an interplay of non-native interactions (See Supporting Information).

## Results

### Computational Folding Time Analysis

The folding time ratios between spectrin domains R16 and R15 (*τ*^*R*16^/*τ*^*R*15^), and R17 and R15 (*τ*^*R*17^/*τ*^*R*15^), for experiments and the different computational models are shown in Figure 1. The first group of bars on the left corresponds to the 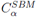 results, in which folding times for R15, R16 and R17 are very similar, *i.e.* ratios close to 1. These results were expected since the three domains are very similar (pair-wise RMSD < 1Å), as has been shown by Best.^4^ The second model, *HP^MJ^*, which takes into account the contribution of non-native hydrophobic interactions, is based on Miyazawa-Jernigan Potential.^25^ The folding times calculated with *HP^MJ^* show a marginal increase compared with those calculated with 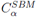, and the relative folding time for R17 increases more than for R16. The third model, *Elec*, the contribution of non-native electrostatic interactions to be taken into account, as described in the Methods section. In comparison with 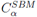 results, *τ*^*R*16^/*τ*^*R*15^ folding time ratio increases, while *τ*^*R*17^/*τ*^*R*15^ decreases, contrary to what happened with the *HP^MJ^* model. The fourth model uses the combination of *HP^MJ^* with *Elec* to perform the folding time simulations. In this case, the folding times of R16 and R17 with *HP^MJ^*+*Elec* are one order of magnitude slower than those of the R15 domain. As for the experimental results, the folding times of R16 and R17 are about three orders of magnitude slower than those of the R15 domain. Even though the results obtained with the last model are distant from those obtained with the experimental one, there is nonetheless an improved qualitative agreement. One can not expect to observe a strict quantitative agreement between computational and experimental folding times, and the difference of two orders of magnitude between experimental and computational results may be understood to be due to the level of simplification of the *C_*α*_* model. Even though the coarse-grained computational model can not be directly and quantitatively compared to the experimental results, it might be able to capture qualitatively the essential features of the folding process of the spectrin domains.

**Figure 1.**
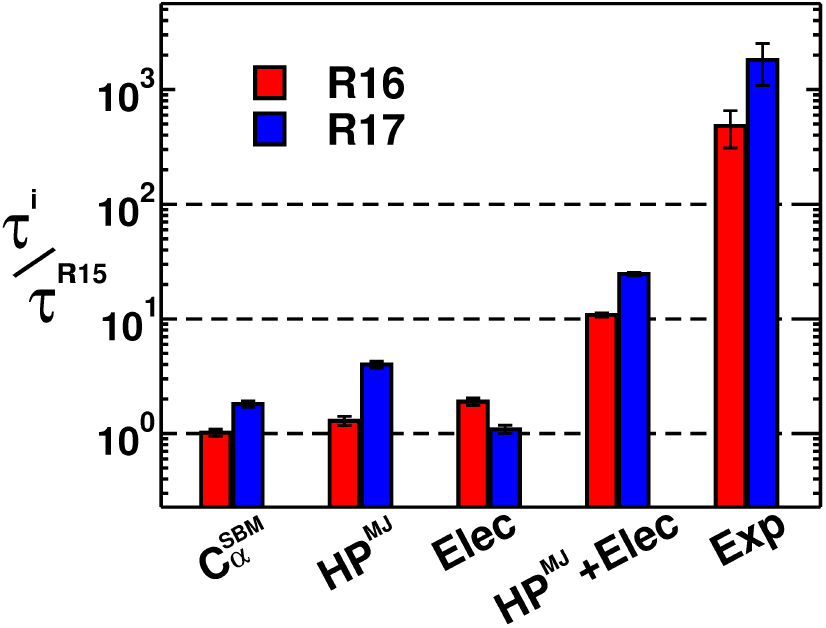
R16 and R17 folding times *τ^i^* normalized by R15 folding time *τ*^*R*15^ for different computational models. The red and blue bars represent the normalized folding times for R16 and R17, respectively. The dashed line equal 1 represents the R15 folding time as a reference. The 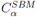 indicates the folding time calculated with the standard SBM model. The *HP^MJ^*and *Elea* represent the folding time simulations with the addition of non-native hydrophobic and electrostatic potential, respectively. *HP^MJ^*+*Elea* is the result for the simulations with both potentials. The last bars represent the folding time from experimental data taken from Wensley et al.^10^

### Mutation Effects in Computational Folding Times

A more comprehensive investigation of the effects of non-native interactions includes simulations with mutations in R16 and R17 domains. The studied mutations were aimed at substituting polar-charged residues with hydrophobic ones and comparing the computational results with experiments.^10,18^ Folding time results performed with *HP^MJ^*+*Elec* potentials are presented in Figure 2A and 2B for R16 and R17, respectively. Figure Figure 2A shows the folding times of R16 normalized by the R15 one for each set of mutations, where M_2_ refers to the double mutation (K25V+E18F) and M_5_ represents the R16 with five mutations (E18F+E19D+I22L+K25V+V30L), and WT is the wild type. All performed mutations are located in the A helix of the spectrin domains, in which residues present in R15 are inserted into R16 and R17.^10^ The computational result presented in Figure 2A shows that R16 with the mutation K25V folds faster than the R16 wild-type. The same occurs for the set of mutations E18F, M_2_ and M_5_, showing that the insertion of R15 residues into R16 speeds up the folding process as also observed experimentally.^10^ The suggestion is that the set of mutations inserting R15 residues into the R16 domain reduces the frustration associated with the search for the correct docking of the helices subsequently reducing the landscape roughness.^10^ The folding times of the R17 domain with the mutations are similar to the folding times of the R16 domain, as is shown in Figure 2B. The R17 *α*-spectrin domains with the set of mutations, K25V, E18F, M_2_ and M_5_ present a faster folding process than the R17 wild-type. The analysis is similar to the R16 domain. The insertion of R15 residues into the A helix of R17 makes this *α*-spectrin domain fold faster, reducing the frustration of the A and C helices interaction.^12,21^

A more extensive set of mutations, which includes mutations that slow down the folding process of R16 and R17 domains,^18,21^ was also investigated computationally. The other mutations in the R16 domain were: H58A, V65A, L87A and A101G.^18^ The mutations performed in the R17 domain were: H58A, V65A, M87A and A100G.^21^ The computational folding times of the R16 and R17 domains, with mutations that speed up or slow down the folding process, were compared with experimental folding times, and are shown in Figures 2C and 2D for R16 and R17, respectively. The computational folding times are in good agreement with the experiments, presenting a significant correlation between these data. The linear correlation for the R16 domain is *R* = 0:94 and for the R17 domain *R* = 0:81. The correlation between the computational and the experimental folding times suggests that the combination of the hydrophobic and the electrostatic potentials can help in the understanding of the basic interactions involved in the protein internal friction phenomenon.

**Figure 2.**
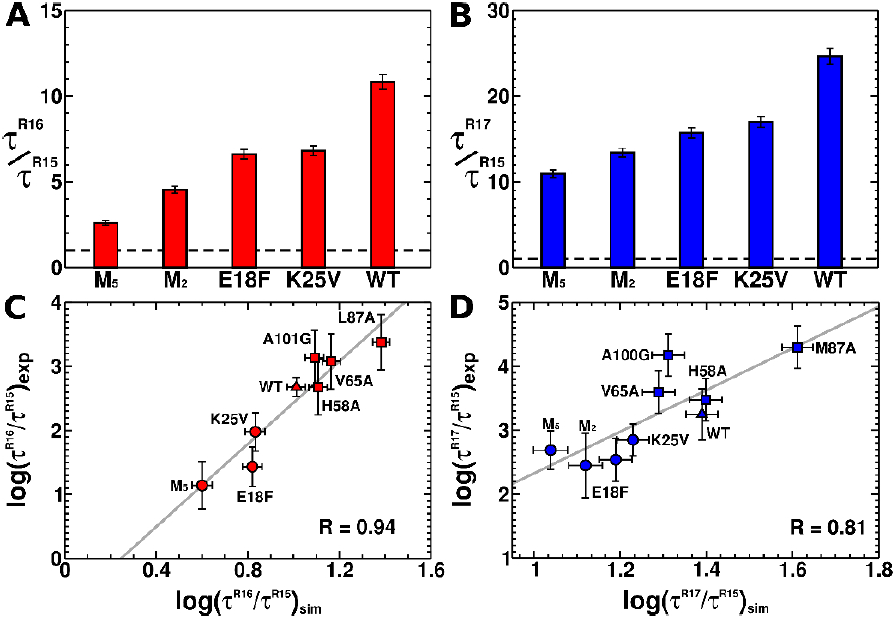
The mutation effects in the folding times for the R16 and R17 domains, *τ*^*R*16^ and *τ*^*R*17^, normalized by the folding time of R15 wild-type,*τ*^*R*15^. (A) The red bars show the folding time for the R16 wild-type (WT) and the mutants K25V, E18F, M_2_, and M_5_. (B) The blue bars show the folding time for the R17 wild-type (WT) and the mutants K25V, E18F, M_2_, and M_5_. (C) Logarithm of *τ*^*R*16^/*τ*^*R*15^ calculated by simulation versus the experimental results, with their respective errors. (D) Logarithm of *τ*^*R*17^=*τ*^*R*15^ calculated by simulation versus the experimental ones, with their errors. (C and D) The circles represent the mutations E18F, K25V, M_2_, and M_5_; the triangles represent the wild-type domains; the squares represent the mutations H58A, V65A, M87A, A100G, and A101G. The solid line represents the fit linear. Experimental data for WT, E18F, K25V, M_2_, and M_5_ for both domains are taken from; Wensley et al. ^10^ R16 mutants H58A, V65A, L87A, and A101G are taken from;Scott et al. ^18^ R17 mutants H58A, V65A, M87A, and A100G are taken from. Scott et al.^21^

## Methods

### Structure-Based C_*α*_ Model

In the Structure-Based Model (SBM), the residues of proteins are represented by individual beads centered on an *α* carbon position.^26–29^ The energy of the protein is given by a Hamiltonian based on its native structure.^26,30^ The energy in a given configuration Γ with regard to the configuration of the native structure Γ_*o*_ is given by

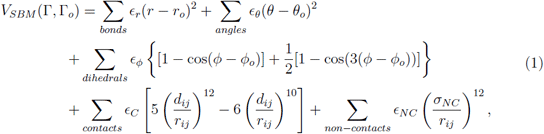

where the distance between two subsequent residues, the angles formed by three and four subsequent residues of native structure are represented by *r*, *θ*, and *ϕ*. The strength of the bonds, angles and dihedral angles is described by *є_r_*, *є_θ_*, and *є_ϕ_*, respectively and the parameter *є_r_* = 100*_є_C__*, *є_θ_* = 20*_є_C__*, *є_ϕ_* = *є_C_*, *є_NC_* = *є_C_* which *є_C_* is equal 1 units (in reduced units). *r_ij_* represents the distance between two non-covalent beads. The interaction of the non-bonded residues in the native state is given by the Lennard-Jonnes 10-12 potential All residue pairs which are not in contact in the native structure interact via non-specific repulsion.

### Nonnative Interactions

#### Heterogeneous Hydrophobic Interactions

This non-native interaction model takes into account the hydrophobic interactions in protein folding. A pairwise hydrophobic amino acid interaction via an attractive Gaussian potential^31–33^ is defined by:

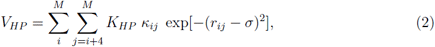

where *M* is the number of hydrophobic amino acids, *r_ij_* is the distance between two hydrophobic amino acids *i* and *j* during the simulation, and *K_HP_* is the overall strength of the hydrophobic forces, with *K_HP_* = 0.1. In the present study, the hydrophobic amino acids considered in this model are Ala, Val, Leu, Ile, Met, Trp, and Phe. The contact energies between two nonnative hydrophobic amino acids *i* and *j* are given by the term *κ_ij_*, with *κ_ij_* = Δ*є_ij_*, where Δ*є_ij_* is the corresponding value from the upper triangle in Table V of Miyazawa and Jernigan^25^ and *σ* = 5.0 Å. The total potential is given by the sum of the SBM potential, *V_SBM_*, plus hydrophobic interactions, *V_HP_*.

#### Electrostatic Interactions

The standard SBM, also know as the vanilla model, does not take into account the charge of the residues explicitly. The electrostatic interactions are explicitly considered by adding charged points at beads, which represent the acidic/basic residues (*i.e.,* histidine, lysine and arginine are positively charged; glutamic acid and aspartic acid are negatively charged). The electrostatic potential, *V_Elec_*, was represented by the Debye–Hückel (DH) model and the interaction between charged residues is given by:

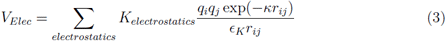

where the charged residues *i* and *j* are represented by *q_i_* and *q_j_*, respectively, *K_electrostatic_*= 332 kcalÅ/(mol *e*^2^), dielectric constant is *є_K_* = 80, *r_ij_* is the distance between charged residues *i* and *j*, and *κ* is the inverse of Debye length.^34^ Therefore, the total potential function of our model is the SBM potential, *V_SBM_*, plus the electrostatic potential, *V_Elec_*.^35–37^

### Simulation Details

All the simulations in this paper were performed using the molecular dynamic package Gro-macs^38^ version 4.5.5 with a leapfrog integration. The input files were obtained with the SMOG@ctbp webtool.^39^ The Berendsen thermostat algorithm^40^ was employed to maintain coupling to an external bath with a constant equal to 1 ps. Proteins were initialized in an open random configuration and simulated over 5×10^9^ steps with time-steps equal to 0.5 fs. The configurations were saved every 5000 steps. The reaction coordinate used to follow the folding events is defined as the fraction of native contacts (*Q*). One native contact between two residues *i* and *j* is considered to have been formed when the distance between them is shorter than 1.2*d_ij_*. The distance *d_ij_* for two residues in the native structure was determined by the software Shadow Contact Map.^41^ The Mean First Passage Time (MFPT) calculations were performed at a different temperature, with each simulation being initialized in an open random configuration (*Q_unf_* ≈ 0.1). The simulation was performed until it reached the folded state, namely, when 80% of native contacts were formed (*Q_fold_* ≈ 0.8). The first passage times were recorded and the MFPT is an average over 100 independent simulations for each temperature. The thermodynamic free energy profile was obtained combining multiple simulations performed over a range of constant temperature runs using the Weighted Histogram Analysis Method (WHAM).^42^ The folding route calculation^43?^ was performed for R15, R16, and R17 in the folding temperature of the R15 domain. The R15 and R16 were cut from residues 1665 to 1771 and 1772 to 1878, respectively, of the full length PDB ID: 1Q4U.^24^ The R17 was cut from residues 115 to 219 of full-length PDB ID: 1CUN.^44^ The mutations in R16 and R17 were generated using Modeller software version 9.17.^45^

### Concluding Remarks

Internal friction terminology has been widely used experimentally as a possible explanation for the difference of three orders of magnitude in folding times between the R15, R16 and R17 domains of *α*-spectrin.^10–12^ In simulations, there are important results in good agreement with experiments with regard to Φ-values and solvent viscosity dependence,^4–6^ but the reason for the folding time differences is still unclear. Best^4^ suggested that the SBM simulations are unable to distinguish the energy landscape roughness of the different *α*-spectrin domains and such roughness must arise from non-native interactions. In the present study, the folding of the R15, R16 and R17 domains of *α*-spectrin using the SBM with the addition of different non-native interaction potentials was investigated. The computational results for folding times obtained from the simulations with the 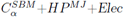 present a reasonable agreement with the experiments. It is possible that the origin of the interactions of the internal friction in *α*-spectrin is due to a cooperativity between non-native hydrophobic and electrostatic interactions. The simulations with each of these potentials separately do not show any agreement with the experimental results. Possibly these two potentials increase frustration and thus the roughness of the energy landscape in R16 and R17, but not in the R15 domain. The difference in the folding kinetic due to the addition of non-native interactions has been reported;^46,47^ an addition of frustration may help the protein to fold faster or slower, depending on the amount of frustration and energy barriers.^28,29^

The effects of addition of different non-native potentials were explored through simulations of folding times for several mutations, in which residues from R15 were inserted into the R16 and R17 domains and then compared with experimental results. A significant correlation between the computational and experimental results was observed.With this evidence, the conclusion is that the origin of the internal friction in *α*-spectrin is related to a combination of non-native interactions, with both hydrophobic and electrostatic contributions. The non-native interactions act in a different way for each *α*-spectrin domain, showing that they are sequence specific. It is perhaps surprising that such a coarse-grained potential can provide a major insight into this complex and long-standing problem. This result may serve as motivation to use the combined *HP^MJ^* + *E_lec_* potential to address other proteins, and serves to demonstrate that the importance of non-native interactions must be taken into account.

## Acknowledgments

We Thank Carolina ATF Mendonça for helpful contributions. FBS was supported by the Higher Education Personnel Improvement Coordination (CAPES) and National Council for Scientific and Technological Development (CNPq - Grant Process No. 141715/2017-0). VGC was funded by Grant 2016/13998-8 and 2017/09662-7, São Paulo Research Foundation (FAPESP) and Higher Education Personnel Improvement Coordination (CAPES). VMO was supported by the National Council for Scientific and Technological Development (CNPq - Grant Process No. 141985/2013-5). VBPL was supported by the National Council for Scientific and Technological Development (CNPq) and FAPESP Grant 2014/06862-7 and 2016/19766-1. We also thank the Center for Scientific Computing (NCC/GridUNESP) of São Paulo State University (UNESP) for computational resources. This work was supported by the Wellcome Trust (grant number WT095195). JC is a Wellcome Trust Senior Research Fellow.

